# Junctophilin-4 is essential for signalling at plasma membrane-endoplasmic reticulum junctions in sensory neurons

**DOI:** 10.1101/842476

**Authors:** Alexandra Hogea, Shihab Shah, Frederick Jones, Chase M Carver, Han Hao, Ce Liang, Dongyang Huang, Xiaona Du, Nikita Gamper

## Abstract

Junctions of endoplasmic reticulum and plasma membrane (ER-PM junctions) serve as signaling hubs in prokaryotic cells. ER-PM junctions are present in peripheral sensory neurons and are necessary for pro-inflammatory G protein coupled receptor signalling and for inflammatory pain generation. Yet, the principles of ER-PM junctions assembly and maintenance, as well as their role in inflammatory signaling in sensory neurons are only beginning to emerge. Here we discovered that a member of the junctophilin family of proteins, JPH4, is abundantly expressed in rat dorsal root ganglion (DRG) neurons and is necessary for the formation of store operated Ca^2+^ entry (SOCE) complex at the ER-PM junctions in response to the G-protein induced ER Ca^2+^ store depletion. Furthermore, we demonstrate a key role of the JPH4 and ER Ca^2+^ stores in the maintenance of inflammatory pain. Indeed, knockdown of JPH4 expression in DRG *in vivo* significantly reduced the duration of pain produced by inflammatory mediator bradykinin. Since the ER supplies Ca^2+^ for the excitatory action of multiple inflammatory mediators, we suggest that junctional Ca^2+^ signalling maintained by JPH4 is an important contributor to the inflammatory pain mechanisms.

## Introduction

Ca^2+^ signals orchestrate processes as diverse as secretion, contractility, excitability, gene expression and cell death. Every mammalian cell expresses a multitude of Ca^2+^-sensitive proteins which have distinct, sometimes even opposing, functional roles. This imposes a requirement for intracellular Ca^2+^ signalling to be spatiotemporally restricted. Such a requirement is particularly evident in mammalian somatosensory neurons; peripheral stimulation excites these neurons locally by activating their Ca^2+^ permeable sensory ion channels or G protein coupled receptors (GPCR) linked to Ca^2+^ signalling pathways. These neurons then generate and transmit action potentials linked to well-defined sensations (touch, temperature, pain, itch etc.). Yet, each neuron hosts a large variety of distinct Ca^2+^ entry and release pathways, thus, the intracellular Ca^2+^ signal fidelity is a key to faithful and specific somatosensory signal transduction.

In order to achieve spatiotemporal segregation of Ca^2+^ signals, the components of the Ca^2+^ signalling cascades are frequently organized into the so-called Ca^2+^ microdomains, specialised regions within a cell, where Ca^2+^ dynamics are locally modulated. These are also regions where Ca^2+^ channels and relevant signalling molecules reside (Berridge, 2006; Parekh, 2008). One example of such microdomains is Ca^2+^ signalling hubs assembled at the junctions between the plasma membrane and the adjacent endoplasmic reticulum (ER-PM junctions) (Berridge, 2006; Pacheco et al, 2016). These junctions are abundant in eukaryotic cells and are increasingly recognized as intracellular signalling ‘hubs’ hosting multiprotein signalling complexes involved in localised GPCR signalling (Jin et al, 2016; Stefan et al, 2013). PM components of these junctional complexes are often located in lipid rafts (Jin et al, 2016; Jin et al, 2013; Pacheco et al, 2016) and include GPCRs that induce Ca^2+^ release from the ER stores via activation of phospholipase C (PLC), generation of inositol 1,4,5-thiophosphate (IP_3_) and subsequent activation of the IP_3_ receptors (IP_3_Rs) localised at the ER side of the junction (Jin et al, 2016; Jin et al, 2013; Stefan et al, 2013). In pain-sensing (nociceptive) sensory neurons, this junctional microdomain signalling is involved in generation of inflammatory pain. Indeed, the inflammatory response of a damaged or infected tissue is orchestrated by a wide range of inflammatory mediators, released into the extracellular space by damaged tissue, recruited immune cells and sensory fibre endings. Many such inflammatory mediators, e.g. bradykinin (BK), proteases, histamine, prostaglandins etc., are capable of exciting nociceptors via their PLC-coupled receptors (reviewed in (Linley et al, 2010; Petho & Reeh, 2012)). Bradykinin receptor-2 (B_2_R) and protease-activated receptor-2 (PAR2) were shown to localise to ER-PM junctions in sensory neurons (Jin et al, 2013). BK is a ‘prototypic’ inflammatory mediator with robust neuronal and vascular action; it is produced at the site of tissue damage/inflammation by the kinin–kallikrein system (Petho & Reeh, 2012). Algogenic activity of BK is so high that it has been branded “the most potent endogenous algogenic substance known” (Dray & Perkins, 1993). B_2_R excite peripheral nerves via several mechanisms, including sensitization of thermo-sensitive TRPV1 channels, inhibition of anti-excitatory M-type K^+^ channels and activation of Ca^2+^-activated Cl^−^ channels (CaCC) (reviewed in (Brown & Passmore, 2010; Jin et al, 2016; Petho & Reeh, 2012)). The latter two excitatory mechanisms are mediated, at least in part, by the ER Ca^2+^ release (Liu et al, 2010; Petho & Reeh, 2012).

In order for PLC-coupled GPCRs to be able to induce and maintain Ca^2+^ release from the ER, the ER needs to be supplied with Ca^2+^ and in most cells this is achieved by the concerted action of a store-operated Ca^2+^ entry (SOCE) complex, delivering Ca^2+^ in to the cytosolic volume of the ER-PM junction and the ER-localised sarcoendoplasmic reticulum Ca^2+^ ATPase (SERCA) (reviewed in (Qiu & Lewis, 2019; Taylor & Machaca, 2019)). In mammalian cells the main components of the SOCE complex are the STIM1-2 proteins, which are the ER Ca^2+^ sensors, and the Orai1-3 proteins, which are the pore-forming subunits of the PM’s Ca^2+^ release activated Ca^2+^ channel, CRAC (Qiu & Lewis, 2019). STIM1 and Orai1 are believed to be the most typical constituents of the SOCE complex in vertebrates. In unstimulated cells Orai1 and STIM1 reside independently in the PM and ER, respectively, but Ca^2+^ release from the ER (e.g. after the GPCR-induced IP_3_R activation) induces STIM1 oligomerization at ER-PM junction, where it binds to and opens Orai1 channels (Qiu & Lewis, 2019; Taylor & Machaca, 2019).

It is still not clear how the ER-PM junctions are formed, maintained and regulated in cells but some proteins localised between the ER and PM membranes were identified and are presumably involved in ER-PM junction assembly and maintenance. These include junctin, junctate, mitsugumin, sarcalumenin, junctophilins and some others (Correll et al, 2017; Hadad et al, 1999; Srikanth et al, 2012; Takeshima et al, 2015).

Molecular components of the SOCE complex have been found in dorsal root-ganglia (DRG) sensory neurons (Gemes et al, 2011; Munoz & Hu, 2016). Previous studies showed that CRAC channels are fully functional in DRG neurons and that *I*_CRAC_ is enhanced after nerve injury (Gemes et al, 2011). STIM1, 2 and Orai1, 3 but not Orai2 were identified in DRG neurons and, interestingly, Orai1 and 3 activation was shown to increase excitability in these cells (Wei et al, 2017).

We hypothesized that SOCE is necessary for maintenance of the inflammatory and proalgesic GPCR signalling at the ER-PM junctions of nociceptive sensory neurons. Indeed, without this functional Ca^2+^ store replenishment mechanism, the persistent Ca^2+^ signalling downstream of pro-inflammatory GPCR activation would not be possible. We further hypothesized that a structural maintenance of the ER-PM junctions is necessary for functional SOCE complex formation. Here we tested these hypotheses using fluorescent imaging, *in situ* proteomics, genetic and pharmacological manipulations and behavioural tests. We discovered that Junctophilin-4 (JPH4) is necessary for the ER Ca^2+^ store refill in DRG neurons. Knock-down of JPH4 in DRG impaired SOCE and weakened the recurrent GPCR signalling *in vitro* and significantly shortened the duration of inflammatory pain *in vivo*.

## Results

### SOCE in peripheral sensory neurons

In order to assess the importance of SOCE for the ER-PM junctional Ca^2+^ signalling and for inflammatory pain generation, we first set to establish a measurement of the SOCE in cultured, small-diameter (mostly nociceptive) DRG neurons. We performed ratiometric Ca^2+^ imaging on DRG neurons loaded with fura-2AM; SOCE was induced with 250 nM BK in Ca^2+^- free extracellular solution for 4 minutes, followed by re-addition of 2 mM Ca^2+^ to the extracellular solution to facilitate *I*_CRAC_. To confirm the identity of the Ca^2+^ transient induced by the extracellular Ca^2+^ add-back we used two SOCE inhibitors: a widely-used pyrazole derivative, YM58483 (BTP2) (Ishikawa et al, 2003), and a newer, selective inhibitor Synta66 (Tian et al, 2016) (**Fig. 1A**). Either YM58483 (1 μM) or Synta66 (3 μM) were applied 1 hour prior to calcium imaging, during fura-2AM loading and were also present throughout the recording. Only neurons responsive to BK were included in the analysis; the BK-responsive DRG neurons are largely TRPV1-positive polymodal nociceptors (Liu et al, 2010). Neither YM58483 nor Synta66 altered the Ca^2+^ release from the ER pool induced by BK, however both drugs almost entirely abolished Ca^2+^ influx during the Ca^2+^ add-back phase of the experiment (**Fig. 1B**). YM58483 and Synta66 reduced the amplitude of the Ca^2+^ add-back transient by 93 ± 44.3% (p<0.001; n=104, N=6) and by 85 ± 12.5% (p<0.001; n=71, N=6), respectively.

**Figure 1.**
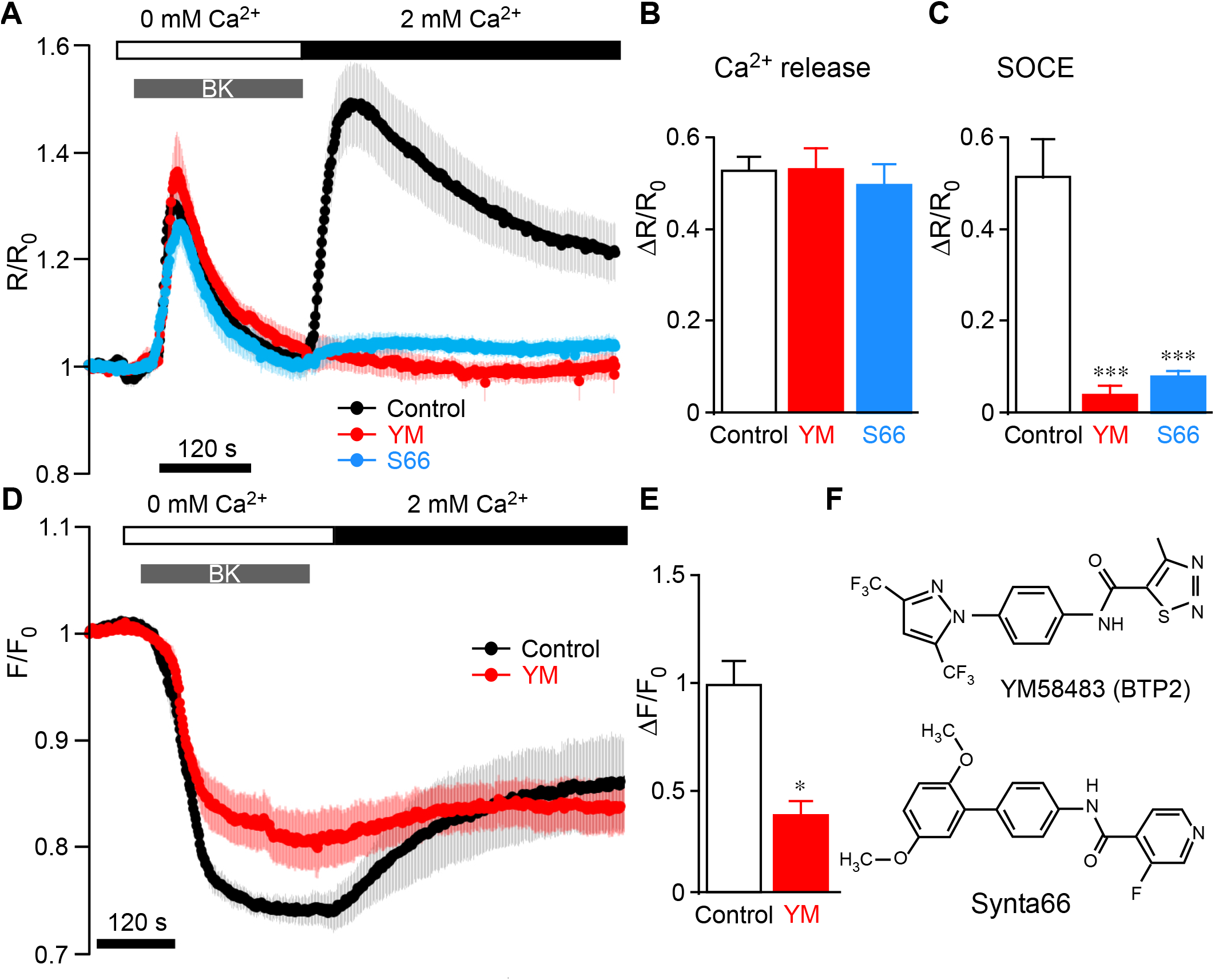
Bradykinin-induced ER Ca^2+^ release and store-operated Ca^2+^ entry (SOCE) in DRG neurons. **A**, Calcium imaging traces of the BK-induced (250 nM) store depletion and Ca^2+^ add-back-induced SOCE in DRG neurons of control neurons (black) and neurons pre-treated with 1 μM YM58483 (red) or 3 μM Synta66 (red) for one hour; the inhibitors were also present extracellular solutions throughout the experiment. Shown is mean normalised F_340_/F_380_ fluorescence intensity ratio (R/R_0_ ± SEM; n = 99 neurons (control), 104 neurons (YM58483) and 71 neurons (Synta66); N=6 independent experiments for each group). Bar charts summarize the effect of YM58483 and Synta66 on the peak BK-induced Ca^2+^ release (**B**) and Ca^2+^ add-back-induced SOCE (**C**); ***, p≤0.001 compared with Control group by one-way ANOVA. **D**, Fluorescent imaging of G-CEPIA1er-transfected DRG neurons in control conditions (black), and after the 1 hr pre-treatment with YM58483 (red); shown is mean normalised fluorescence (F/F_0_ ± SEM; n=3 for each group). E, bar chart summarizing the effect of YM58483 on the BK-induced ER Ca^2+^ depletion; * p<0.05, unpaired Student’s t-test. (**F**) Chemical structure of SOCE inhibitors YM58483 and Synta66, used in this study.

In order to confirm that BK indeed induces ER store depletion and that Ca^2+^ add-back results in ER store refill due to SOCE, we transfected acutely dissociated DRG neurons with an ER-localised calcium-measuring organelle-entrapped protein indicator (G-CEPIA1*er*)(Suzuki et al. 2014). DRG neurons displayed robust ER Ca^2+^ depletion in response to BK, which was partially recovered during the Ca^2+^ add-back (**Fig. 1C**). Treatment with YM58483 (1 μM) significantly reduced ER Ca^2+^ uptake during the Ca^2+^ add-back to 37±9% of the levels seen in the control group (p = 0.04, n=3; **Fig. 1C**). The amplitude of the BK-induced drop in G-CEPIA1*er* fluorescence was somewhat smaller after YM58483 treatment but the difference did not reach significance. Due to low efficiency of transfection of DRG neurons with G-CEPIA1*er*, only a small number of individual experiments were successfully completed. The experiments presented in **Fig. 1** confirmed that BK induces both the ER Ca^2+^ release and SOCE, with the latter being sensitive to SOCE/CRAC inhibitors, YM58483 and Synta66.

We next thought to confirm presence in DRG neurons of the major proteins constituting SOCE complex: STIM1-2 and Orai1-3. Immunohistochemical staining of DRG sections demonstrated that all STIM and Orai isoforms, except of Orai2, are expressed in DRG neurons (**Fig. 2**). Orai1 was expressed in 72% of all neurons tested (1263/1774); immunoreactivity had obvious PM localization (**Fig. 2A**, inset). The average somatic diameter of the Orai1-positive cells was 21.5 ± 0.9 μm, very similar to the mean soma size of all DRG neurons analysed (21.4 ± 1 μm) hence, there was no evident preference of Orai1 expression in neurons of a particular size, suggesting broad expression among neurons of different sensory modalities (**Fig. 2A**, left, **D**). Co-staining with neuronal markers NF200 (myelinated Aβ and Aδ neurons; **Fig. S1A**) and TRPV1 (mostly polymodal nociceptors; **Fig. S1B**) confirmed ubiquitous distribution of Orai1 in DRG. Orai2 antibody signal was localised to the nucleus (**Fig. 2A**, middle) and was therefore not analysed. Orai3 immunoreactivity was observed in 51% (304/594) of all DRG neurons analysed (**Fig. 2A**, right, **E**); Orai3 immunoreactivity was detected with significantly lower incidence as compared to Orai1 (p<0.01; Fisher’s exact test). The mean somatic diameter of the Orai3-positive neurons was 22.67 ± 1.69 μm, suggesting expression across the neurons of different sensory modalities.

**Figure 2.**
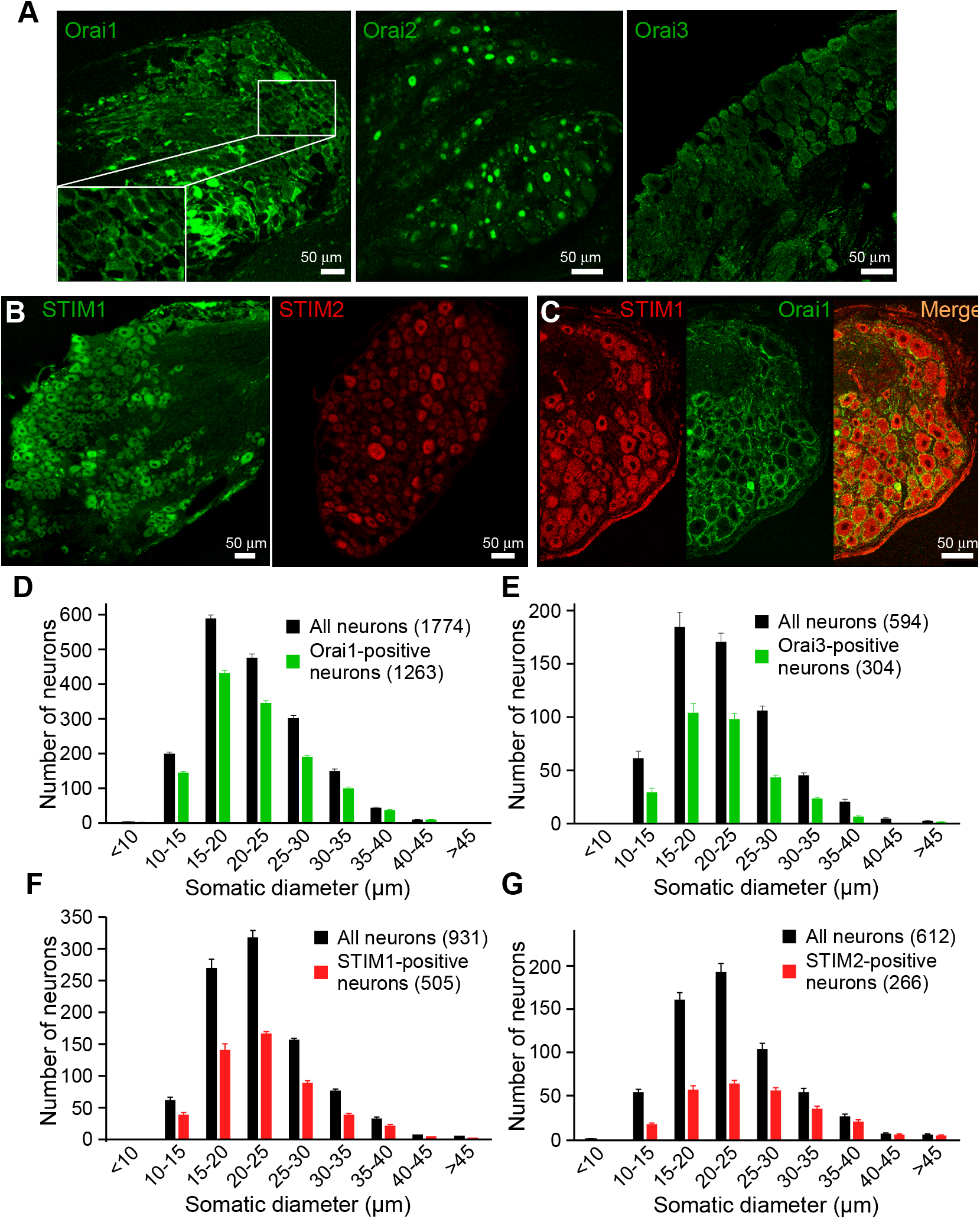
Expression of Orai and STIM isoforms in the DRG. Confocal micrographs for immunostaining of DRG sections with antibodies against Orai1-3 (**A**), STIM1 and STIM2 (**B**) and co-immunostaining of Orai1 and STIM1 (**C**). Scale bars = 50 μm. **D-G**, The somatic diameters of neurons positive for Orai1 (**D**), Orai3 (**E**), STIM1 (**F**) or STIM2 (**G**) were measured and plotted as a frequency histograms alongside the total number of neurons (black). Data are shown as the number of immunofluorescence-positive neurons within each somatic size-band ± SEM; cumulative data from 10 individual DRG sections for Orai1 and from 5 individual sections for the other proteins shown.

STIM1 was expressed in 54% (505/931) of all neurons analysed; the mean somatic diameter of the STIM1-positive cells was 23.3 ± 1.2 μm, similar to the average soma size of all DRG neurons analysed in these sections (23.9 ± 1.5 μm; **Fig. 1B**, left, **F**). Co-staining with NF200 (**Fig. S1C**) and TRPV1 (**Fig. S1D**) confirmed that STIM1 is expressed in medium and large myelinated neurons, as well as in TRPV1-positive nociceptors. STIM2 immunostainings demonstrated that 43% (266/612) of all DRG neurons analysed contained this protein (**Fig. 2B**, right, **G**), an incidence which was significantly lower as compared to that of STIM1 (p<0.01; Fisher’s exact test). The mean somatic diameter of STIM2- positive neurons was 25.2 ± 2.6 μm, slightly greater than the overall average of 22.8 ± 2 μm, suggesting higher expression in larger neurons.

Because Orai1 and STIM1 were the most abundant subunits of those tested and because Orai2 was found not necessary for SOCE in DRG neurons (Wei et al, 2017), we concluded that the ‘classic’ Orai1-STIM1 complex is the most abundant SOCE correlate in the majority of DRG neurons. We therefore focused on this complex in subsequent experiments. Contribution of other subunits to SOCE in DRG neurons however cannot be excluded.

### Expression of junctophilins in sensory neurons

ER-PM junctions are maintained by junctional protein anchors and of these, the junctophilin family is perhaps one of the best understood (Takeshima et al, 2015). Moreover, deletion or downregulation of JPH1 and JPH2 has been shown to impair SOCE in muscle cells (Hirata et al, 2006; Li et al, 2010), while downregulation of JPH4 impaired SOCE in T-cells (Woo et al, 2016). Yet, to the best of our knowledge the expression profile and potential function of juctophilins in sensory neurons is hitherto unreported. Thus, we performed immunohistochemistry and western blot experiments to investigate expression of juctophilins in DRG (**Fig. 3**); both methods revealed no detectable levels of JPH2 (**Fig. 3A, E**). On the other hand, expression of JPH1, JPH3 and JPH4 was found in 52% (456/882), 38% (302/798) and 76% (805/1066) of DRG analysed, respectively. JPH4 was expressed in a significantly higher proportion of neurons, as compared to JPH1 and JPH3 (p<0.01, Fisher’s exact test). Neuron size distribution analysis of JPH1, 3 and 4 expression did not reveal a bias towards neurons of a particular size (**Fig. 1B-D**). NF200 and TRPV1 co-localization experiments demonstrated that JPH4 was highly co-expressed with these neuronal markers; 75 of the NF200-positive and 84% of the TRPV1-positive neurons also displayed JPH4 immunoreactivity (**Fig. S2**). Western blot analysis (**Fig. 3E**) confirmed presence of JPH1, 3 and 4, but not JPH2 in DRG while all JPH proteins were detected in their specific control tissues. Interestingly, the JPH4 antibody revealed a double band in DRG lysate, one of which was of the predicted molecular weight (66 kDa), yet, only a lighter band was detected in the brain sample, suggesting existence of at least two forms of this protein (e.g. alternatively spliced gene products or a post-transcriptional modification). Taken together, the above experiments identified JPH4 as a major junctophilin in the DRG.

**Figure 3.**
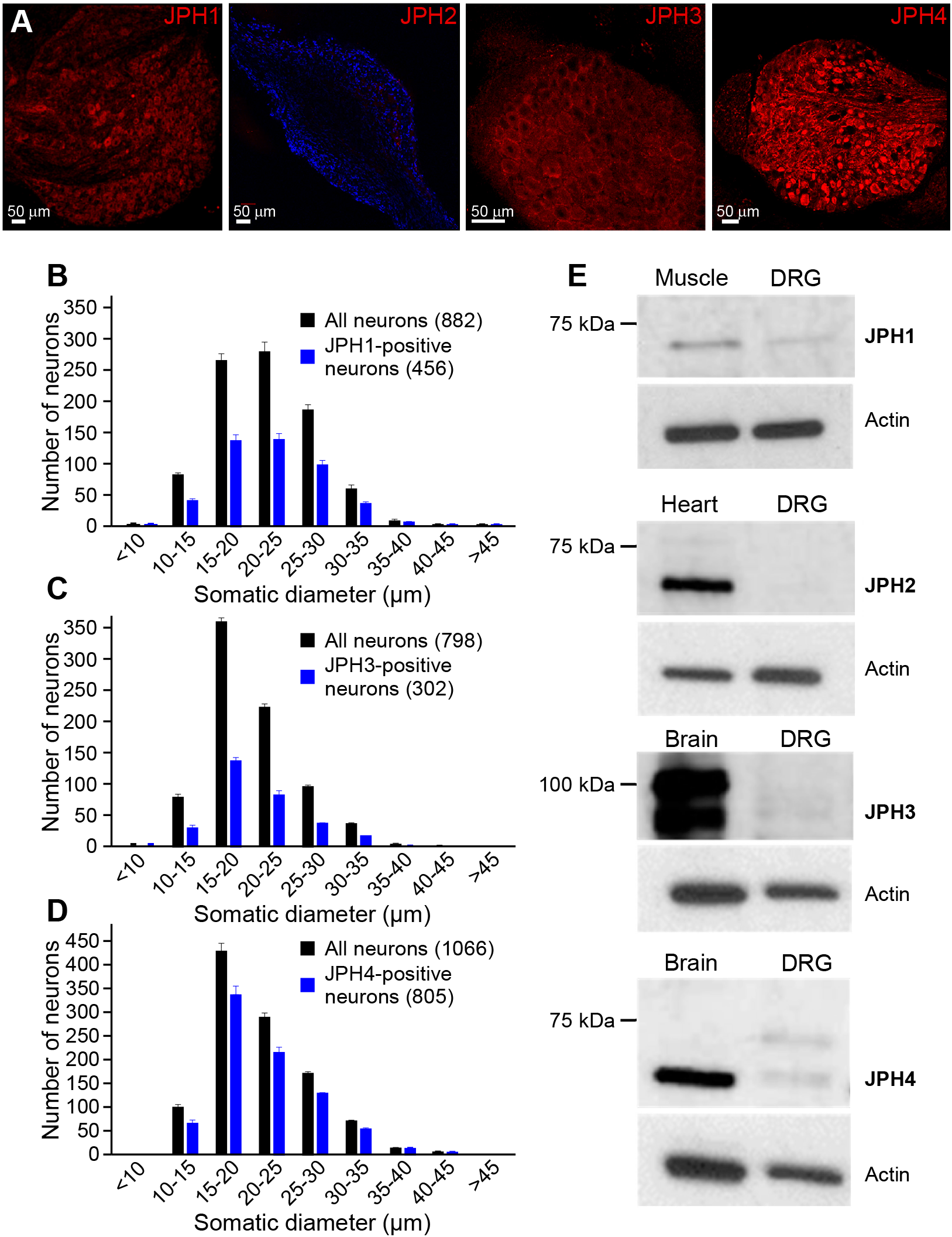
Expression of Junctophilin isoforms in the DRG. **A**, Confocal micrographs for immunostaining of DRG sections with antibodies against JPH1-4, as indicated, DAPI staining is included in the JPH2 micrograph to depict DRG section. Scale bars = 50 μm. **B-D**, The somatic diameters of neurons positive for JPH1 (**B**), JPH3 (**C**) or JPH4 (**D**) were measured and plotted as a frequency histograms alongside the total number of neurons (black). Data are shown as the number of immunofluorescence-positive neurons within each somatic size-band ± SEM; cumulative data from 6 individual DRG sections for each protein analysed. **E**, Western blot analyses of the JPH family members in DRG and positive control tissue lysates (Skeletal muscle for JPH1, Heart for JPH2 and Brain for JPH3 and 4). These are representative results of 6 individual experiments for each JPH isoform.

Co-staining of JPH4 with either Orai1 (**Fig. 4A**) or STIM1 (**Fig. 4C**) revealed a high degree of co-expression of these proteins in neurons. **Fig. 4B** and **Fig. 4D** display small-diameter DRG neurons in culture imaged using an improved-resolution Airyscan method (LSM 880, Zeiss). Orai1, JPH4 and STIM1 all displayed fluorescence maxima at the PM; the Orai1/JPH4 (**Fig. 4B**) and STIM1/JPH4 (**Fig. 4D**) PM maxima significantly overlapped, suggesting close proximity at the ER-PM junctions. In many individual DRG neuron somata JPH4 immunoreactivity displayed striking tubular appearance with a preferred distribution towards the PM, rather than being randomly diffused in the cytoplasm; an example of such JPH4-positive tubular network is given in **Fig. 4E**. These observations suggest that JPH4 might be present at the ER-PM junctions of DRG neurons.

**Figure 4.**
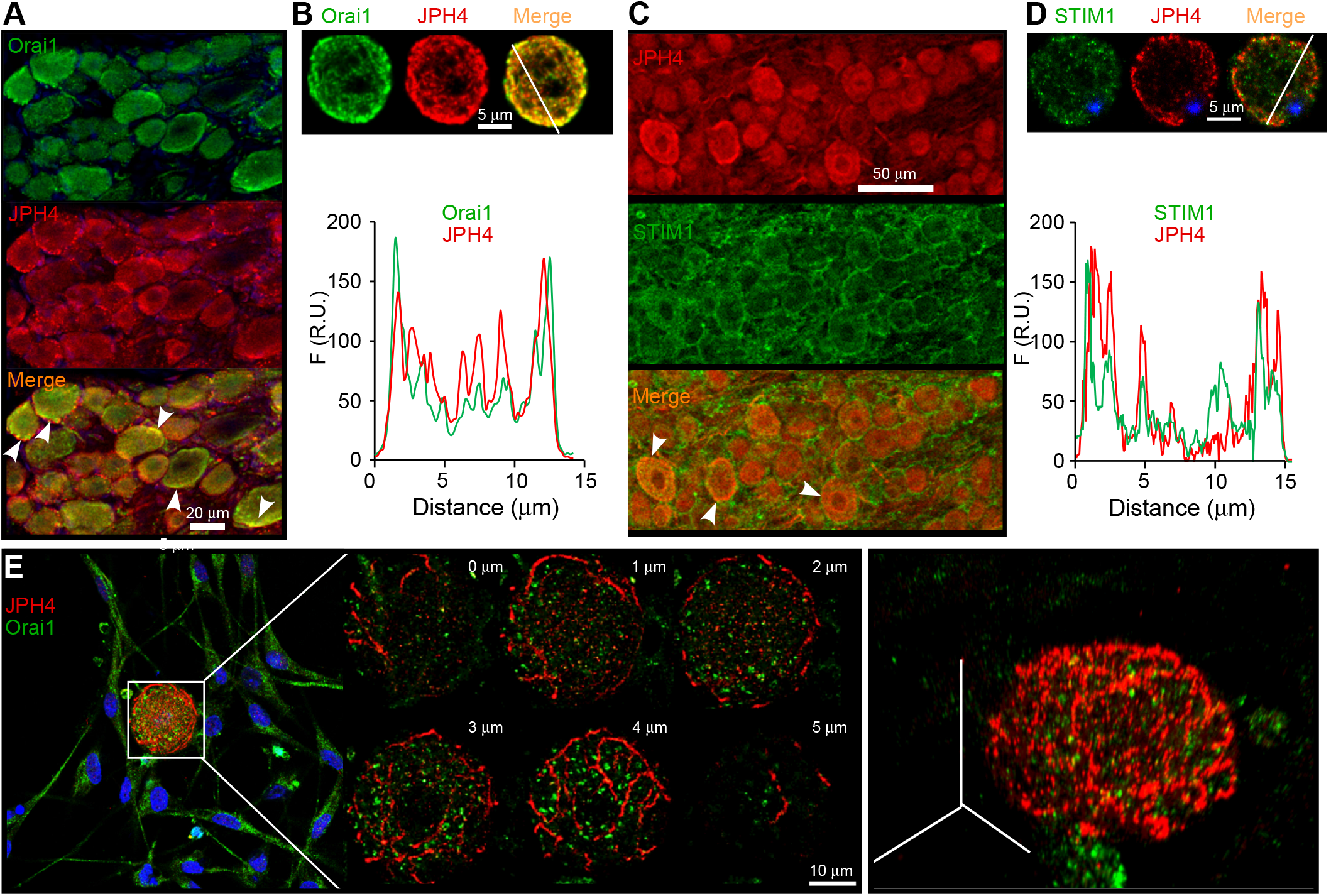
Intracellular localization of JPH4 in DRG neuron somata. **A, B**, Co-immunostaining of Orai1 (green) and JPH4 (red) in DRG. **A**, co-staining of PFA-fixed DRG sections. **B**, Airyscan images of co-labelled DRG neuron in culture, lower panel shows the fluorescence intensity line scan displaying overlapping intensity maxima for both proteins at the PM. **C, D**, Co-immunostaining of STIM1 (green) and JPH4 (red) in DRG. **C**, co-staining of PFA-fixed DRG sections. **D**, Airyscan images of co-labelled DRG neuron in culture, lower panel shows the fluorescence intensity line scan displaying overlapping intensity maxima at the PM and some additional intracellularly localized maxima. **E**, Airyscan image (Z-stack of 1 μm optical sections) of a cultured DRG neuron revealing a specific tubular localization of JPH4 (red) near the plasma membrane. Right panel is a 3D representation of the same neuron.

### Role of JPH4 in junctional signal transduction in DRG neurons

Since clustering of STIM1 with Orai1 and SOCE complex formation is a signature response to GPCR-induced ER store depletion (Qiu & Lewis, 2019), we used proximity ligation assays and co-immunoprecipitation to test if JPH4 is important for this process in DRG. Immunoprecipitates from DRG neurons endogenously expressing JPH4 and STIM1 revealed a weak interaction between the two proteins at rest (**Fig. 5A**). BK treatment resulted in increased co-immunoprecipitation signal, suggesting increased interaction between JPH4 and STIM1 during the BK-induced ER store depletion (**Fig. 5A**). To confirm this interaction, we used an ‘in situ proteomic’ approach, proximity ligation assay (PLA). This method allows specific detection of closely associated proteins *in situ*. Only if the proteins of interest are within no more than 30-40 nm proximity, bright fluorescent PLA puncta of 0.5-1 μm diameter are formed ((Soderberg et al, 2006; Weibrecht et al, 2010); see Methods). Proximity between STIM1 and JPH4 (**Fig. 5B, C**), Orai1 and JPH4 (**Fig. 5D, E**) and Orai1 and STIM1 (**Fig 5F, G**) was significantly increased following the BK treatment (250 nM; 15 min in Ca^2+^-free extracellular solution). This suggests that while some level of clustering between JPH4, STIM1 and Orai1 might already exist at resting conditions, the clustering is significantly increased in response to the ER Ca^2+^ release. This, in turn, might suggest that JPH4 facilitates SOCE by either promoting ER-PM proximity or by directly facilitating STIM1-Orai1 interactions. Specificity of PLA has been confirmed by either omittiong one of the primary antibodies, while keeping both secondary PLA probes (negative control; **Fig. S3A**) or by using two different primary antibodies against the same protein (positive control, **Fig. S3B**).

**Figure 5.**
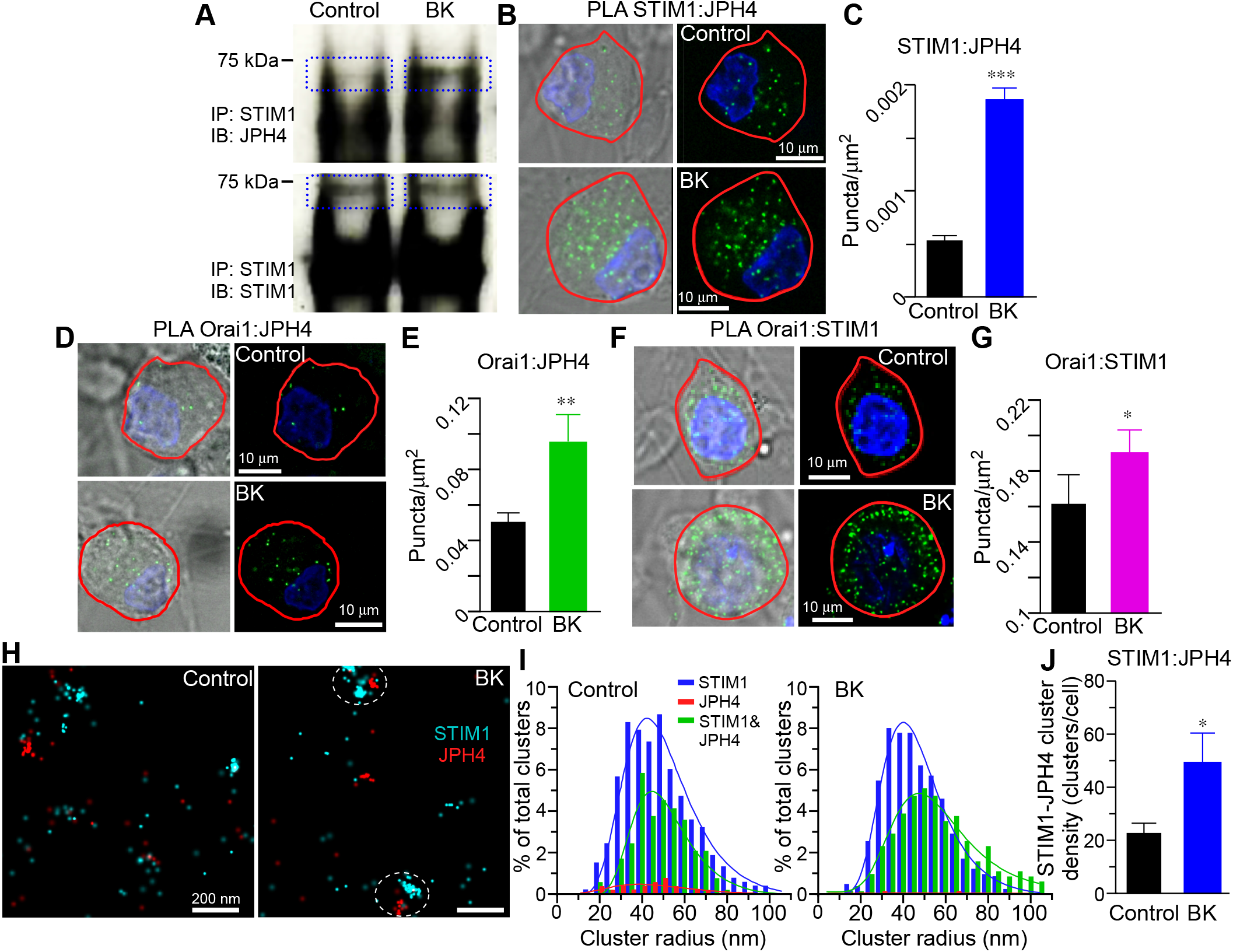
JPH4 interacts with the SOCE complex in DRG neurons. **A**, JPH4 co-immunoprecipitates with STIM1. Immunoprecipitation from DRG neurons was performed with the anti-STIM1 antibody and immunoblotted for the detection of JPH4 and STIM1. Cells were left untreated (Control) or treated with 250 nM BK for 15 min in Ca^2+^ free solution to induce store-depletion before lysis; representative of 6 independent experiments. **B**, Proximity ligation assay (PLA) probing JPH4 and STIM1 proximity in small diameter DRG neurons. Cells were left untreated (Control) or treated with 250 nM BK for 15 min in Ca^2+^ free solution to induce store-depletion. **C**, Quantification of PLA puncta density in experiments as those shown in panel B; n = 24 neurons (control), 14 neurons (BK); N=6 independent experiments. Scale bars = 10 μm. **D-G** Experiments similar to that shown in panels **B, C** but for Orai1:JPH4 (n = 60 neurons (control), 34 neurons (BK); N = 6 independent experiments) and Orai1:STIM1 (n=33 neurons, N=6 individual experiments) proximity. Labelling and experimental conditions as in B and C. **H**, representative STORM images from control or BK-treated (250 nM BK for 15 min in Ca^2+^ free solution) DRG neurons, double-labelled for STIM1 using dye-pairs of AF405/647 (cyan centroids) and JPH4 using dye-pairs of Cy3/647 (red centroids); scale bars = 200 nm. **I**, Comparison of distributions of clusters representing double-labelled STIM1-JPH4 under control and BK-stimulation conditions, respectively (see Methods for detail). **J**, Quantification of cluster density in experiments as those shown in panels H, I; n = 7 neurons (control), 8 neurons (BK). In Panels C, E, G and J data shown as mean ± SEM, ***p<0.001; ** p<0.01; *p<0.05; Student’s t-test.

To further confirm the co-localization of STIM1 and JPH4 after BK-induced ER store depletion, we performed super-resolution STORM imaging (Zhang et al, 2016). Control or BK-treated (250 nM; 15 min in Ca^2+^-free extracellular solution) DRG cultures were fixed and labelled with primary antibodies for STIM1 and JPH4, followed staining by in-house secondary antibodies conjugated to photoswitchable fluorophores (see Methods). Using near-TIRF, oblique-angle illumination, nanodomain proximity of STIM1 and JPH4 has been observed, denoted by clustered STORM co-localizations (**Fig. 5H, I**). In the BK-treated neurons, there were significantly more detected STIM1- JPH4 clusters compared to control (**Fig. 5J**), consistent with the PLA and Co-immunoprecipitation experiments. In examination of the population distribution of detected clusters, there was both a significantly increased percentage of STIM1-JPH4 interactions (**Fig. 5J**) and right-shift in cluster size (**Fig. 5I**) in BK-treated DRGs compared to control, but there were no significant differences between conditions in the STIM1-only or JPH4-only clusters.

Next, we tested if JPH4 knockdown in DRG neurons would affect STIM1-Orai1 interaction. Two different siRNA targeting JPH4 were used at various concentrations, separately or in combination to optimize the protocol. The siRNA2 (see methods) demonstrated the highest efficacy of reducing JPH4 protein expression in DRG neurons 48 hours post transfection, as tested using immunohistochemistry and western blot analysis (**Fig. 6A, B**). PLA experiments testing the interaction between Orai1 and STIM1 were performed on DRG neurons transfected with JPH4 siRNA2 or scrambled oligo. Treatment of DRG neurons transfected with scrambled oligo with BK (250 nM; 15 min in Ca^2+^-free extracellular solution) significantly increased Orai1-STIM1 interaction, revealed as PLA puncta, as compared to vehicle treated control (Fig. 6C, E). This was consistent with experiments on untransfected cells (**Fig 5D, G**). Strikingly, no significant difference between the number of puncta in BK-treated and vehicle-treated neurons was detected when JPH4 was silenced (Fig. **6D, F**). These experiments strongly suggest that JPH4 is necessary for the SOCE complex formation in DRG neurons.

**Figure 6.**
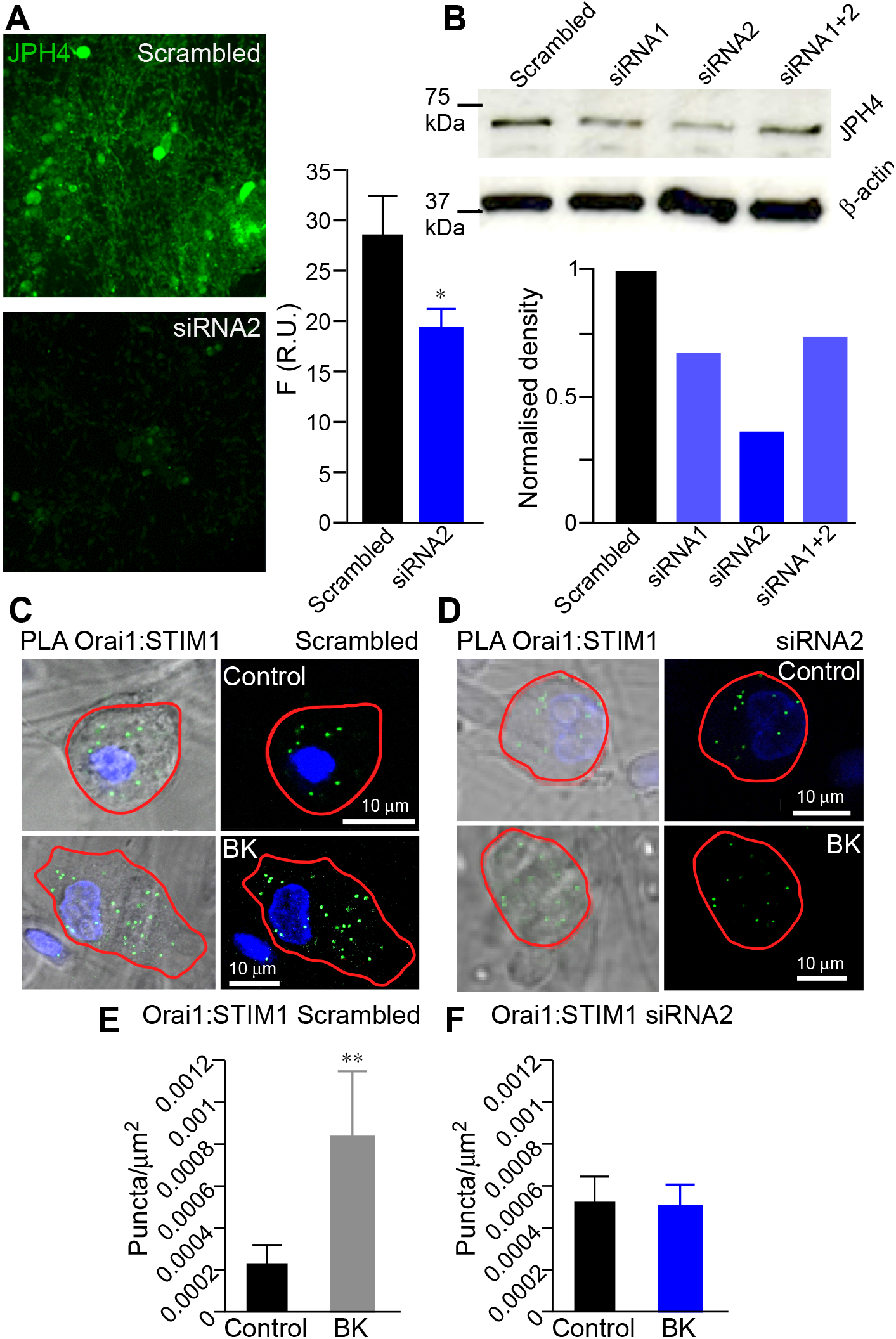
Knockdown of JPH4 in DRG neurons disrupts STIM1:Orai1 clustering. **A**. Assay of JPH4 immunofluorescence in DRG culture transfected with scrambled oligo or JPH4-siRNA2. The data are summarised in the bar chart on the right; n=6 for both data sets; *p<0.05; Student’s t-test. **B**, Western blot analysis of JPH4 abundance in DRG lysates from cultures transfected with JPH4-siRNA1, JPH4-siRNA2 or scrambled oligo. **C, D**, Representative example of fluorescent PLA signals between Orai1 and STIM1 in cultured DRG neurons transfected with scrambled oligo (**C**) or JPH4-siRNA2 (**D**). Cells were left untreated (scrambled oligo: n = 10, N = 5; JPH4 siRNA2: n = 9, N = 6) or treated with 250 nM BK for 15 min in Ca^2+^ free solution (scrambled oligo: n = 6, N = 6; JPH4 siRNA2: n = 9, N = 6) to induce store-depletion. **E, F**, Quantification of PLA puncta density in experiments as those shown in panels **C-D**. Data shown as mean ± SEM, ** p<0.01; Student’s t-test.

Next, we set out to test how JPH4 knockdown would affect BK-induced Ca^2+^ signalling in DRG neurons. In the first series of experiments we compared the amplitudes of BK-induced and Ca^2+^ add-back induced Ca^2+^ transients in DRG neurons transfected with either JPH4 siRNA2 or scrambled oligo using experimental protocol presented in the **Fig. 1B**. Silencing JPH4 in DRG did not significantly affect the amplitude of the BK-induced Ca^2+^ transient (**Fig. 7A, B**) but reduced the SOCE (Ca^2+^ transient during the Ca^2+^ add-back) by 45 ± 14.3% (n=8; p<0.05, Student’s t-test; **Fig. 7A, C**). These results suggest that JPH4 might be involved in facilitating SOCE; yet, the basal ER Ca^2+^ levels could be maintained in unstimulated cells even with such partial SOCE impairment.

**Figure 7.**
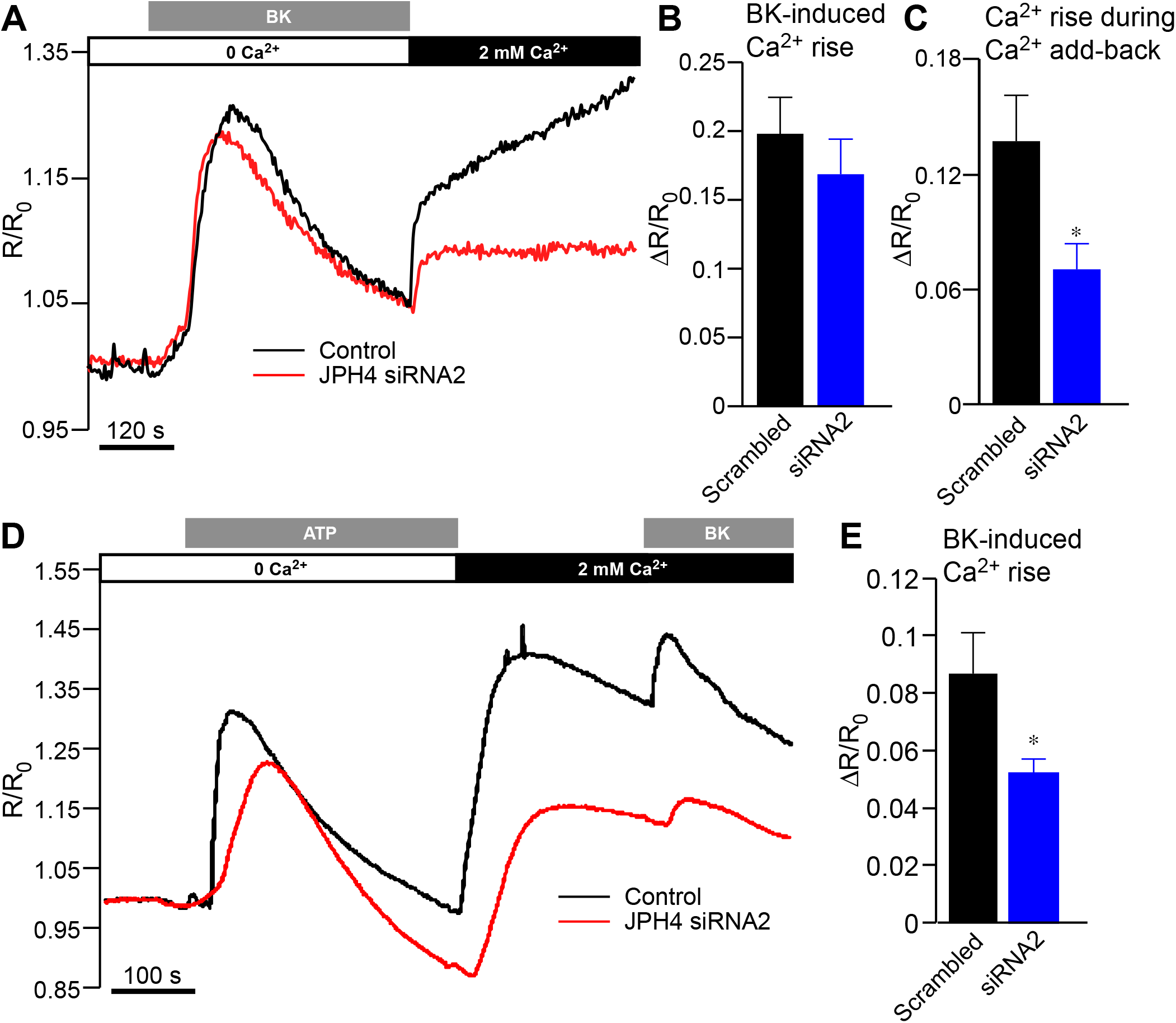
Knockdown of JPH4 in DRG neurons disrupts SOCE and junctional Ca^2+^ signaling. **A**, Representative example of Fura2 calcium imaging recording of typical store depletion/Ca^2+^ add-back experiments in control (scrambled oligo) and JPH4-siRNA2 transfected DRG neurons. **B, C**, Summary data for experiments as these shown in **A**; the effect of JPH4 KD on the amplitude of the BK-induced Ca^2+^ transients (**B**) and SOCE during the Ca^2+^ re-addition phase (**C**). Data are shown as mean ± SEM; scrambled oligo: n = 9, N = 3; JPH4 siRNA2: n = 8, N = 3; *p<0.05 Student’s t test. **D**, Representative example of Fura2 calcium imaging recording in control (scrambled oligo) and JPH4-siRNA2 transfected DRG neurons. First store depletion was induced by ATP (30 μM) in the Ca^2+^ free buffer; Ca^2+^ was then added back to the extracellular solution and secondary IP_3_-dependent store depletion with BK (250 nM) was performed. **E**, Summary data for experiments as these shown in **D**: the effect of JPH4 KD on the amplitude of the BK-induced Ca^2+^ transients during the Ca^2+^ re-addition phase. Data are shown as mean ± SEM; scrambled oligo: n = 6, N = 3; JPH4 siRNA2: n = 6, N = 3 *p<0.05, Student’s t-test.

We then reasoned that while ER Ca^2+^ stores can be maintained in unstimulated neurons even with JPH4 knocked down, impaired SOCE in JPH4-deficcient neurons could impede recurrent GPCR signalling as ER stores would not replenish fully in such conditions. To test this hypothesis we designed an experimental protocol where two rounds of store depletion were applied in sequence. To avoid desensitization and tachyphylaxis common for the GPCR, including B2R (Bawolak et al, 2009), DRG neurons were treated with two different GPCR-agonists. Thus, cultured cells were first stimulated with 30 μm ATP (in Ca^2+^-free extracellular solution) to activate endogenous Gq/11-coupeld P2Y receptors expressed in DRG neurons (Gerevich & Illes, 2004). Following the Ca^2+^ add-back phase, BK was applied to trigger second round of the IP_3_-dependent store depletion, this time by activating B_2_R. It has to be pointed out that even though DRG neurons also express ionotropic ATP-activated P2X_2_/P2X_3_ receptors (Gerevich & Illes, 2004), our experimental protocol excluded the contribution of these ion channels to [Ca^2+^]_i_, since application of ATP was made in Ca^2+^-free extracellular solution (in line with previous protocols). **Fig. 7D, E** shows that the secondary (BK-induced) Ca^2+^ release from the stores was indeed reduced by 40.0 ± 9.8% (n=6; p<0.05, Student’s t-test) in JPH4-silenced neurons as compared to scrambled oligo controls. The ATP-induced Ca^2+^ transients were also somewhat smaller in JPH4-silenced neurons, but this difference did not reach significance. Taken together, the experiments presented in **Fig. 7** demonstrate that *i)* JPH4 knockdown impairs SOCE and *ii)* it reduces the capacity of the ER Ca^2+^ stores to sustain recurrent GPCR-mediated Ca^2+^ signalling.

### Knockdown of JPH4 in vivo shortens the duration of inflammatory nociception

In the next series of experiments we asked if JPH4 knockdown and impaired SOCE would affect the ability of BK to generate pain *in vivo*. Indeed, BK-induced nociception depends, at least in part, on the ER Ca^2+^ release and subsequent activation of excitatory Ca^2+^-activated Cl^−^ channels (CaCC) and suppression of inhibitory M-type K^+^ channels (Lee et al, 2014; Liu et al, 2010; Petho & Reeh, 2012). Thus, we reasoned that since JPH4 knockdown impairs STIM1-Orai1 clustering and inhibits SOCE, it may also affect the ability of BK to sustain inflammatory nociception *in vivo*. To knock-down JPH4 *in vivo* we conjugated JPH4 siRNA2 oligo to cholesterol for *in vivo* RNA interference (Wolfrum et al, 2007) and performed intrathecal injections to the L5/L6 area for DRG delivery (see Methods). Cholesterol-conjugated JPH4 siRNA2 efficiently knocked down JPH4 expression in DRG culture *in vitro* (**Fig. 8A**); intrathecal application of siRNA induced marked downregulation of JPH4 in the whole DRG *in vivo* by 41.3±9.6% (n=6; p<0.01, Kruskal-Wallis ANOVA with Mann-Whitney test). Next, we performed hind-paw injections of BK to rats, which received either intrathecal JPH4 siRNA2, scrambled oligo or a vehicle and analysed pain-related ‘nocifensive’ behaviour (time spent licking, biting and flinching the injected paw). Nocifensive behaviour after the BK injection (10 nmol/site; 50 μl) in either of the control groups lasted for about 15 minutes. We binned the time after injection in 2-min intervals and calculated the mean time of nocifensive response within each 2-min interval (**Fig. 8C**). Interestingly, the initial (peak) response to BK was not affected by the JPH4 knockdown, but the effect tailed off significantly faster in JPH4-knocked-down animals, as compared to either of the controls (**Fig. 8C**). This finding is consistent with our hypothesis that JPH4 silencing compromises the ability of DRG neurons to replenish their ER Ca^2+^ stores, thus making them to desensitize towards the BK-induced excitation much faster. Total duration of nocifensive behaviour over the 15 minutes observation period was also significantly reduced by the JPH4 silencing (**Fig. 8C**). Importantly, general sensitivity of the cutaneous afferents to mechanical or thermal stimulation (as tested by von Frey or Hargreaves tests, respectively, see Methods) was not affected by JPH4 knockdown (**Fig. 8E, F**). In sum, the experiments presented in **Fig. 8** strongly suggest that JPH4 knockdown *in vivo* impairs junctional, GPCR-induced Ca^2+^ signalling in sensory neurons hence accelerating their desensitization in inflammatory environment.

**Figure 8.**
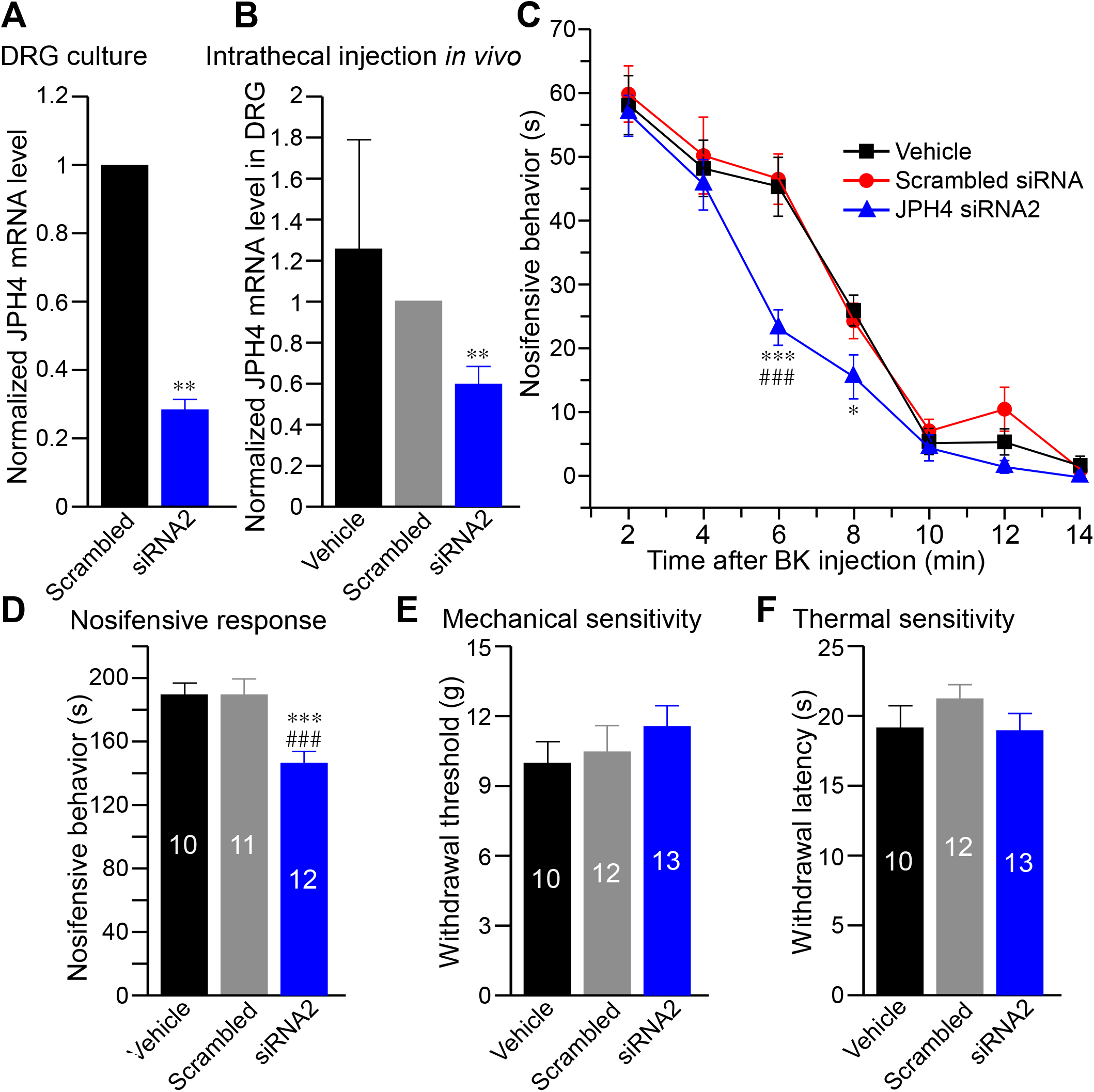
Knockdown of JPH4 in DRG *in vivo* attenuates BK-induced pain. **A, B**, Efficiency of the JPH4 KD using the cholesterol-conjugated JPH4-siRNA2 as tested by RT-PCR. **A**, Application of cholesterol-conjugated JPH4-siRNA2 or scrambled oligo (200 nM) to dissociated DRG neurons in culture; n = 5; **p<0.01, Student’s t-test. **B**, Intrathecal injections (2 pmol/site; twice a day for 4 days) of JPH4-siRNA2, scrambled oligo or vehicle *in vivo*. L5-L6 DRG were analysed for JPH4 expression using RT-PCR. **C**, Intrathecal JPH4 KD attenuated BK-induced nocifensive behaviour. BK (10 nmol/site; 50 μl) was injected into the right hind paw and the time spent licking, biting and flinching the injected paw was recorded and plotted in 2-min intervals. ***p<0.001, *p<0.05 as compared against scrambled oligo; ^###^p<0.001 as compared to vehicle (vehicle: n=10; scrambled oligo: n=11; JPH4-siRNA2: n=12). **D**, total time of nocifencive behaviour over 30 min observation period in the same animals analysed in panel **C**. **E, F**, Effect of JPH4 KD on the mechanical (**E**, withdrawal threshold, von Frey method) and thermal (**F**, withdrawal latency, Hargreaves method) sensitivity (vehicle: n=10; scrambled oligo: n=12; JPH4-siRNA2: n=13).

## Discussion

In the present study we investigated junctional Ca^2+^ signalling in peripheral somatosensory neurons. We discovered that JPH4 is the major junctophilin expressed in this type of neurons and that JPH4 is necessary for the formation of SOCE complex at ER-PM junctions in response to the GPCR-induced ER Ca^2+^ store depletion. Furthermore, we demonstrate a key role of JPH4 and ER Ca^2+^ stores in the maintenance of inflammatory pain *in vivo*. Since the ER supplies Ca^2+^ for the excitatory action of BK and many other inflammatory mediators that are coupled to G_q/11_-type of Gα protein and PLC-mediated signalling cascades (Choi & Hwang, 2018; Linley et al, 2010; Liu et al, 2010; Petho & Reeh, 2012), we suggest that junctional Ca^2+^ signalling maintained by JPH4 is an important contributor to the inflammatory pain mechanisms.

We started by characterization of DRG neuron SOCE, a key Ca^2+^ signalling process that requires ER-PM junctions to occur. We found that in DRG neurons that respond to BK, there is robust SOCE, which is sensitive to *I*_CRAC_ inhibitors, YM58483 (Ishikawa et al, 2003) and Synta66 (Tian et al, 2016). We further show that STIM1-2 and Orai1, 3 are broadly expressed in DRG neurons, with STIM1 and Orai1 being the most abundant subunits expressed in the majority of these neurons; these data are in accord with recent publications (Qi et al, 2016; Wei et al, 2017). Interestingly, while STIM1, Orai1 and Orai3 were present amongst neurons of all sizes, indicating no clear bias towards neurons of specific sensory modality, STIM2 was much more frequently detected in large-diameter neurons: 34% of the neurons with diameter ≤ 25 μm and 80% of neurons with diameter ≥ 40 μm were found to be STIM2-positive (**Fig. 2G**). Thus, STIM2 could have a specific function in myelinated Aβ fibers mostly responsible for different aspects of mechanosensation (Basbaum et al, 2009). Since in this study we were mainly interested in the role of junctional signalling in nociceptors, we focused here on STIM1 and Orai1, which we found to be the most abundant SOCE components in small-diameter, TRPV1 positive nociceptors.

To further characterize junctional signalling in DRG neurons we investigated the presence of junctophilins in DRG. This family of junctional proteins is fairly well characterized in muscle and immune cells, but virtually nothing is known about presence and function of these proteins in sensory neurons.

Proteins of the mammalian JPH family (JPH1-4) have their N- and C-termini residing in the PM and ER, respectively. The N-terminus contains eight ‘membrane occupation and recognition nexus’ (MORN) repeats, which have high affinity for PM phospholipids and act as a ‘PM anchor’, while the C-terminal trans-membrane segment spans the SR/ER membrane (Kakizawa et al, 2007). JPH1 and JPH2 are found in SR-PM junctions of skeletal and cardiac muscle cells, respectively, and facilitate the interaction of PM-localised voltage-gated Ca^2+^ channels and SR-localised Ryanodine receptors, thus, orchestrating excitation-contraction coupling (Takeshima et al, 2015). JPH3 and JPH4 were mostly found in the CNS but their role is much less understood. It was reported that JPH3/JPH4 double knockout mice exhibited irregular hind limb reflexes and impaired memory; it was further suggested that these deficits arose from the impaired functional coupling between the NMDA receptors, ryanodine receptors, and SK K^+^ channels due to compromised ER-PM junctions (Moriguchi et al, 2006). Another recent study reported that JPH3 and JPH4 are necessary for functional coupling of L-type Ca^2+^ channels, ryanodine receptors and SK K^+^ channels in hippocampal neurons to mediate slow afterhyperpolarization (Sahu et al, 2019). Additionally, JPH4 and junctate were found to form a complex that allows STIM1 recruitment after ER Ca^2+^ discharge in T-cells (Woo et al, 2016).

Here we show that JPH4 is abundantly expressed in most (over 75%) DRG neurons, JPH2 is virtually absent whilst JPH1 and JPH3 are present but in a smaller proportion of neurons and their immunoreactivity is also less prominent than that of JPH4. These findings allow us to conclude that JPH4 is the main junctophilin isoform present in DRG. Airyscan imaging revealed a striking pattern of JPH4 localization in tubular structures running along the PM in DRG somata and co-localisation of JPH4 with both STIM1 and Orai1 at or near the PM. These observations suggest that JPH4 may indeed represent an important mechanism for ER-PM junction formation and maintenance in sensory neurons.

Using proximity ligation and STORM we show that in DRG neurons JPH4 indeed resides in close (less than 40 nm) proximity of both STIM1 and Orai1. Interestingly, ER Ca^2+^ store depletion in response to BK not only stimulated clustering of STIM1 and Orai1, as is expected form the SOCE complex formation, but also significantly increased clustering of JPH4 with either STIM1 or Orai1 (**Fig. 5**). Thus, it is reasonable to suggest that JPH4 is an important element of SOCE. In support of this idea, siRNA knockdown of JPH4 abolished BK-induced STIM1-Orai1 clustering (**Fig. 6**) and impaired functional SOCE (as tested with Ca^2+^ imaging; **Fig. 7**).

Using JPH4 knockdown as a tool to compromise the ER Ca^2+^ store refill, we show that JPH4 deficiency significantly reduces the ability of the ER to sustain Ca^2+^ transients during repetitive store depletion (**Fig. 7D**). Moreover, intrathecal knockdown of JPH4 in DRG *in vivo* significantly shortened the duration of the BK induced pain without affecting the basal thermal or mechanical sensitivity of cutaneous fibers (**Fig. 8**). The later notion is an important indication that the JPH4 knockdown specifically affects only these aspects of nociceptive signalling, that depend on junctional Ca^2+^ signalling.

It is important to note that the majority of the data presented here are obtained using DRG neuron cell bodies, but inflammatory excitation of peripheral fibres occurs at the nerve endings and the peripheral sections of the nerves surrounded by the inflamed tissue. Several considerations allow us to hypothesize that the effects observed at the somatic levels are also relevant to the peripheral segments of the somatosensory fibers. Thus, *i)* BK injections are painful, suggesting presence of functional B_2_R at the cutaneous nociceptive terminals (Lee et al, 2014; Liu et al, 2010). *ii)* BK-induced pain, at least in part, depends on the Ca^2+^-induced activation of ANO1 channels, an effect confined to the ER-PM junctions (Jin et al, 2016; Jin et al, 2013; Lee et al, 2014; Liu et al, 2010). *iii)* ER, or ‘axoplasmic reticulum’ is abundantly present in C-fibres; it runs up until the proximal segments of the nociceptive free nerve endings (Kruger et al, 2003) and, finally, *iv)* intrathecal knockdown of JPH4 reduces the duration of BK-induced pain (present study, **Fig. 8**), suggesting the importance of junctional Ca^2+^ signalling at the cutaneous nociceptive terminals for the BK-mediated inflammatory nociception.

In summary, this study identified JPH4 as a crucial element of localised Ca^2+^ signalling at ER-PM junctions in nociceptive sensory neurons and established the importance of such signalling for inflammatory nociception.

## Materials and Methods

All animal work carried out at the University of Leeds was performed under UK Home Office License and in accordance with the regulations of the UK Animals (Scientific Procedures) Act 1986. Animal experiments performed in Hebei Medical University were in accordance with the Animal Care and Ethical Committee of Hebei Medical University, (Shijiazhuang, China) under the International Association for the Study of Pain (IASP) guidelines.

### Cell culture, transfection and siRNA delivery in vitro

Rat DRG neurons were dissociated and cultured as described previously (Jin et al, 2013; Kirton et al, 2013; Liu et al, 2010). In brief, Wistar rats (7 to 14 days old) were sacrificed by isoflurane overdose, followed by cervical dislocation. Rats were decapitated, the spine removed and sectioned longitudinally. DRG were removed and dissociated in HBSS containing 10 mg/ml dispase (Thermo Fisher Scientific) and 1 mg/ml type 1A collagenase (Sigma-Aldrich) in a humidified incubator at 37°C and 5% CO_2_ for 13 minutes. Ganglia were then gently triturated and re-incubated for a further 2 minutes, followed by final trituration and addition of ice-cold DMEM with GlutaMAX, 10% FCS, penicillin 50 U/ml and streptomycin 50 μg/ml. Cells were then washed twice by centrifugation, resuspended in pre-warmed media and plated on 10 mm glass coverslips pre-coated with poly-D-lysine (Millipore Ltd) and laminin (Insight Biotechnology). Cells were then incubated for 4 hours to facilitate adhesion; each coverslip was then supplemented with 1 ml culture media and incubated for a minimum of 48 hours prior to experiments.

Transfection with calcium-measuring organelle-entrapped protein indicator (CEPIA) was performed as previously described (Kirton et al, 2013). Briefly, a Nucleofector® I console (Lonza) was used to transfect DRG neurons. DRG dissociation was performed as described above. Prior to seeding, cells were re-suspended in a rat neuron transfection buffer (Lonza) and mixed with 10 μg of cDNA. The solution was then quickly transferred into a Lonza certified cuvette and placed into the Nucleofector. Program O-03 was utilized for transfection; the cells were then re-suspended in pre-warmed media and seeded onto coverslips. As for standard cultured DRGs, cells were incubated for 48 hours prior to experiments.

For gene silencing experiments, small interfering RNA (siRNA) oligos against JPH4 were transfected into the DRG with Lipofectamine RNAiMAX Transfection Reagent (Thermo Fisher Scientific). DRG neurons were dissociated as previously described; 48 hours post-seeding (approx. 70% confluence) the cultures were washed with fresh pre-warmed DMEM (without serum and antibiotics). The transfection reagent and siRNA were mixed in OptiMEM I reduced serum medium (Life Technologies) as per manufacturer’s instructions. The diluted siRNA-lipid complex was then added to cells followed by incubation at 37°C for 1 to 3 days. Two different siRNA products against JPH4 (Silencer Select, s175513 and s175511, Life technologies) as well as scrambled control (Silencer Select Negative Control No.1, Life Technologies) were used at 10 pmol per well in both single use or in combination.

Human embryonic kidney 293 (HEK 293) cells were cultured on 100 mm culture dishes in Dulbecco’s Modified Eagle’s medium (DMEM-GlutaMAX, Life Technologies) supplemented with 10% Foetal Calf Serum (FCS, Sigma-Aldrich), 50 U/ml penicillin (Sigma-Aldrich) and 50 μg/ml streptomycin (Sigma-Aldrich) and kept in a humidified incubator (set at 37°C and 5% CO2). Cells were passaged every 3 days. HEK293 cells were transfected in the 24 well plates using Fugene HD (Promega) according to the manufacturer’s instructions. Imaging experiments were performed 48 hours post-transfection.

### Fluorescence Ca^2+^ imaging

Plated DRG cultures were loaded with fura-2 by 1-hour incubation with a mixture of 2 μM fura-2 AM (Thermo Fisher Scientific) and 0.01% pluronic acid (Sigma-Aldrich) in extracellular bath solution, EC, of the following composition (in mM): 160 NaCl, 2.5 KCl, 1 MgCl_2_, 2CaCl_2_, 10 HEPES, 10 glucose (pH 7.4). The mixture was then removed and cells were washed with EC solution. Fragments of coverslips were placed on the microscope perfusion chamber and EC solution was perfused over the cells through a gravity driven perfusion system at an approximate rate of 2 ml/min. The compounds were diluted in EC solution to the required concentration and applied via the perfusion system. The Ca^2+^-free EC contained (in mM): 160 NaCl, 2.5 KCl, 1 MgCl_2_, 1 EGTA, 10 HEPES, 10 glucose (pH 7.4). Fluorescence imaging was performed on an inverted Nikon TE-2000 microscope and was connected to a Till Photonics fluorescent imaging system comprising of a Polychrome V monochromator and an IMAGO CCD camera. TILL Vision 4.5.56 or Live Acquisition 2.2.0 (FEI) were used for image acquisition and processing. Cells were excited at 340 nm and 380 nm (50 ms exposure) and the emission was collected using a UV-2A filter (Nikon). Regions of interest (ROIs) were used to select neurons from snapshots taken to record from and for further post-hoc analysis. Recordings made were analysed in Microsoft Excel and Origin software.

Green CEPIA1*er cDNA* was purchased from Addgene. Imaging of ER calcium levels using G-CEPIA1*er* was performed in the same manner as fura-2 imaging however single-wavelength excitation at 488nm was used.

### Immunohistochemistry

DRG were collected, washed with PBS and fixed with 4% ice-cold paraformaldehyde (PFA, Sigma-Aldrich) for 1 hour. The ganglia were then embedded in 10% gelatine (Sigma) solution prepared with distilled water. The gelatine was then cut into 1 cm^3^ cubes containing a single ganglion, and further incubated in 4% PFA for 6 to 10 hours at 4°C. The cubes were washed with PBS, cut to 40 μm sections using a microtome (Leica VT 1200S) and washed three times with PBS. Sections were then incubated with a blocking buffer (containing 0.05% Tween 20, 0.25% Triton X-100 and 5% donkey and/ or goat serum in PBS, all products are from Sigma-Aldrich) for 1 hour at room temperature. Subsequent incubation was done overnight at 4°C with a solution containing one or two primary antibodies (**Table S1**), depending on the experimental design. The antibodies were diluted to the required concentration in 5% bovine serum albumin (BSA, Sigma-Aldrich) in PBS. Next day, sections were washed with PBS and further incubated with fluorochrome-conjugated secondary antibodies (**Table S1**) diluted in 1 % BSA buffer for 2 hours at room temperature. The DRG slices were then mounted with Vectashield plus DAPI, covered with coverslips and sealed with nail polish prior to imaging. Sections were kept in the dark, at 4°C and imaged with a Carl Zeiss LSM880 inverted confocal microscope and processed with Zeiss ZEN imaging software. For each staining procedure, a buffer without primary antibody was used as a negative control and cells were detected by nuclear DAPI staining. Antibodies against STIM, Orai, NF200 and TRPV1 were explicitly characterized in previous studies (**Table S1**). Specificity of JPH antibodies was tested with Western blot in tissue homogenates known to express each of the proteins (**Fig. 3E**). To analyse the expression of STIM, Orai and JPH proteins, the number of positively and negatively stained neurons was determined. The mean fluorescence intensity of the whole DRG section was considered the threshold above which a neuron was considered ‘positive’. The cell diameter and mean intensity was calculated for each selected neuron. Cells were classed depending on the somatic diameter and presented as a frequency distribution. Data are shown as numbers of positive neurons of different somatic sizes from the total number of neurons analysed. Cultured DRG neurons were fixed with 4 % ice-cold PFA on glass coverslips for 20 minutes, the staining procedure was similar to that applied to DRG sections.

### Proximity Ligation Assay

The PLA (Duolink, Sigma-Aldrich) was performed according to the manufacturer’s instructions and as described previously (Jin et al, 2013). In this approach, the proteins of interest within a cell/tissue are labelled with specific primary antibodies and then treated with PLA probes, which are secondary antibodies conjugated with short DNA oligos. If two proteins reside within less than 30-40 nm of each other, a connector oligonucleotides facilitate the formation of a single-stranded DNA circle between the two secondary probes, a unique new DNA sequence is amplified, and a colour reaction is developed. DRG cultures were fixed with 4% ice-cold PFA for 20 minutes and permeabilised with buffer composed of 0.05% Tween 20 and 0.25% Triton X in PBS for 1 hour at room temperature. Cells were then blocked for 30 minutes at 37°C (using Duolink blocking buffer) and incubated with primary antibodies (**Table S1**) for 12-16 hours at 4°C. Following washing, the oligo-conjugated secondary antibodies (anti-mouse MINUS and anti-rabbit PLUS PLA probes, Sigma-Aldrich) were added for 1 hour at 37°C. Ligation and amplification was performed according to the manufacturer’s instructions. Finally, the samples were washed and mounted with Duolink mounting medium with DAPI (Sigma-Aldrich). Imaging was performed using a Carl Zeiss LSM880 inverted confocal microscope and images processed by Zeiss ZEN software.

### Stochastic Optical Reconstruction Microscopy (STORM)

The multicolour STORM was performed as described by one of us recently, using the same set of equipment and controls (Zhang et al, 2016). Briefly, images were acquired on a Nikon N-STORM super-resolution system (Nikon Instruments Inc.), consisting of a Nikon Eclipse Ti inverted microscope and an astigmatic 3D lens placed in front of the EMCCD camera to allow the Z coordinates to be most accurately determined. Two-color STORM laser control was performed with non-overlapping activator dyes, Alexa 405 carboxylic acid (Invitrogen, #A30000) and Cy3 mono-reactive dye pack (GE Healthcare, #PA23001), conjugated to affinity-purified secondary antibodies from Jackson Immunoresearch, along with the reporter dye, Alexa Fluor 647 carboxylic acid (Invitrogen, #A20006). The extent of co-localization of STIM1 and JPH4 was determined with the same primary antibodies as in PLA experiments (**Table S1**). The activator and reporter fluorophores were conjugated in-house to an appropriate unlabeled secondary antibody. STORM imaging was performed in a freshly prepared imaging buffer that contained (in mM): 50 Tris (pH 8.0), 10 NaCl and 10% (w/v) glucose, with an oxygen-scavenging GLOX solution (0.5mg/ml glucose oxidase (Sigma-Aldrich), 40μg/ml catalase (Sigma-Aldrich), and 10mM cysteamine MEA (Sigma-Aldrich). MEA was prepared fresh as a 1M stock solution in water. Acquisitions were made from between 2-3 different experiments for labelling and imaging. Images were rendered as a 2D Gaussian fits of each localization. The diameter of each point is representative of the localization precision (larger diameter, less precise), as is intensity (more intense, more precise). Signal-noise thresholds were handled as peak height above the local background in the N-STORM software Elements). A detected peak was set as the central pixel in a 5×5 pixel area, and the average intensity of the 4 corner pixels was subtracted from intensity of the central pixel. Using a 100x objective and 16×16μm pixel are of the iXon3 camera, this corresponds to a 0.8×0.8μm physical neighbourhood.

### Cluster size and localization proximity analysis

Unfiltered STORM localization data were exported as molecular list text files from Nikon-Elements and were analysed with in-house software incorporating a density-based spatial clustering of applications with noise (DBSCAN) (Zhang et al., 2016). A dense region or cluster was defined as localizations within a directly-reachable radius proximity (epsilon) from a criterion minimum number of other core localizations (MinPts). Density-reachable points were localizations that were within the epsilon radius of a single core point and thus considered part of the cluster. Localizations considered to be noise were points that were not within the epsilon distance of any core points of a cluster. We derived the appropriate epsilon parameter using the nearest-neighbour plot from single-dye labelled controls; cluster detection was determined for epsilon between 20 and 80nm, which were the nearest-neighbour localization distances representing 95% of area under the curve. DBSCAN parameters were verified by measuring goodness of fit to Gaussian distribution with cluster population data from single-dye labelled controls, and set for distance of directly reachable points at 50nm (epsilon) and 5 minimum points (MinPts). These parameters were found to be the most stringent possible, that also reliably fit the control data. For cluster detection, each localization was assessed based on its corrected X and corrected Y 2D spatial coordinates, and the associated activator dye was tracked throughout analysis. Detected clusters were tabulated by the composition of resident activator dyes contributing to the total neighborhood of localizations for that cluster. Clusters were categorized based on activator dye composition as Alexa405 only, Cy3 only, or Alexa405 + Cy3 according to the dye conjugated to each antibody label. Cluster radius size (nm) data were placed in probability distribution histograms with bin size of 5nm. Based on the observation that a majority of cluster distributions displayed positive skew, distributions were fit by the generalized extreme value distribution function: 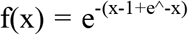. Cluster size values were reported as mean ± SEM. In addition, the percentage of clusters belonging to a labelling category out of total clusters was compared. Population statistics for each cluster category were derived from 7-8 cells per staining and imaging condition.

### Western blot and immunoprecipitation

DRG neurons from two 7 days old Wistar rats were seeded on 6-well plates. Cells were lysed 48 hours post-seeding with lysis buffer (in mM): 50 HEPES, 150 NaCl 183.9 Na_3_VO_4_, 10 NaF, EDTA with 10% Glycerol and 1% NP-40; supplemented with protease inhibitor. Lysates were homogenised and centrifuged for 10 minutes at 13,000 g at 4°C. DRG lysates were boiled for 5 min in SDS–polyacrylamide gel electrophoresis (SDS-PAGE) sample buffer (50 mM tris HCl (pH 6.8), containing 5% 2-mercaptoethanol, 10% glycerol, and 1% SDS) and analysed by SDS-PAGE, followed by transfer onto a polyvinylidene difluoride membrane by electroblotting. The membranes were incubated in blocking buffer (tris-buffered saline, TBS, containing 5% skimmed milk powder and 0.1% Tween 20 for 1 hour, followed by incubation with primary antibodies (**Table S1**), diluted in the same buffer, at 4°C overnight. The membranes were washed in TBS containing 0.1% Tween 20 before 45 min incubation with an appropriate secondary antibody (horseradish peroxidase– conjugated anti-rabbit or anti-mouse secondary antibody). The bands were visualized using an enhanced chemiluminescence substrate (Pierce ECL WB Substrate and Super Signal West Femto Thermo-Scientific). All samples were re-probed for β-actin as a loading control. The exposed films were processed using a Xograph machine, scanned and analysed using ImageJ software.

For co-immunoprecipitation DRG lysates were prepared as described above. A pre-cleaning step was included by adding 8 μl of beads (Protein A.G PLUS-Agarose, Santa Cruz Biotechnology) to 500 μg lysate and mixing for 30 minutes at 4°C to reduce non-specific binding. The mixture was then centrifuged at 9,000 g for 5 minutes at 4°C. The supernatant was collected and incubated with the appropriate primary antibody (Table S1) for 2 hours at room temperature. On the following day, antibody-coated protein G–Sepharose beads (GE Healthcare; 20 μl) were added to the mixture and incubated on a shaker overnight at 4°C. The beads were washed with PBS 3 times, centrifuged and supernatant discarded. SDS-PAGE was performed as described above.

### In vivo siRNA knockdown

The *in vivo* JPH4 knock-down was performed as described earlier (Luo et al, 2005) by using the JPH4 siRNA2 oligonucleotide conjugated to cholesterol (conjugation was performed by RiboBio). Cholesterol-conjugated JPH4 siRNA2 or scrambled control oligo were reconstituted in saline. The siRNA (200 nM, 10 μl) or vehicle were injected intrathecally (i.t.) between the L5 and L6 dorsal spinous processes as described earlier (Huang et al, 2016; Zhang et al, 2019) under isoflurane anaesthesia. The treatment was repeated twice daily for 4 days. The pain tests were performed on the next day after the last injection. After behavioral testing, animals were humanly sacrificed; lumbar DRGs were extracted and analysed for the knockdown efficiency by RT-PCR.

### Behavioral assays

Adult rats (body weight, 170–180 g) were allowed to acclimate for at least 30 min in a transparent observation chamber before the experiment. For BK-induced pain assessment, animals received a 50-μl intraplantar injection of BK (10 nmol/site) or saline into the right hind paw. Animals were video-recorded for 30 minutes after the injection and protective (‘‘nocifensive’’) behavior was analysed as time spent licking, biting, lifting, and flinching the injected paw. The analyses were performed by an observer who was unaware of treatment allocations. For measurement of thermal sensitivity, the change in latency of hind-paw withdrawal in response to noxious heat was recorded using the Hargreaves’ plantar method (Ugo Basile); the heat source was set to 20% of the maximal intensity. Sensitivity to mechanical stimuli was assessed with von Frey method, as described previously (Chaplan et al, 1994). Briefly, calibrated nylon filaments (von Frey hair, Stoelting) with different bending forces were applied to the midplantar surface of the right hindpaw. The filaments were applied starting with the softest and continuing in ascending order of stiffness. A brisk withdrawal of the right hind limb was considered a positive response. Each stimulus was applied for 5 times with an interval of 5 seconds. The threshold was recorded when 3 or more clear withdrawal responses within a set of 5 applications were observed with a given filament.

## Acknowledgements

We thank Eva Jaworska and Stephen Milne for expert technical assistance. This work was supported by the BBSRC grants BB/R003068/1 and BB/R02104X/1 to N.G., Wellcome Trust Investigator Award 212302/Z/18/Z to N.G.; National Natural Science Foundation of China grants 81701113 to D.H. and 31571088 to X.D.; the Excellent Youth Fund of Natural Science Foundation of Hebei (C2018206222) to D.H.; the Key Basic Research Project of Applied Basic Research Program of Hebei Province (16967712D) to X.D.

## Author contributions

A.H. planed, performed and analysed experiments, S.S. assessed the effects of JPH4 knockdown using PLA, analysed data; F.J. performed IHC analysis of JPH expression, C.M.C. performed STORM experiments; H.H., C.L., D.H. and X.D. performed *in vivo* siRNA experiments and behavioural tests; N.G. designed the study, analysed data and wrote the manuscript (assisted by all the co-authors).

## References

Basbaum AI, Bautista DM, Scherrer G, Julius D (2009) Cellular and molecular mechanisms of pain. Cell 139: 267–284

Bawolak MT, Fortin S, Bouthillier J, Adam A, Gera L, R CG, Marceau F (2009) Effects of inactivation-resistant agonists on the signalling, desensitization and down-regulation of bradykinin B(2) receptors. Br J Pharmacol 158: 1375–1386

Berridge MJ (2006) Calcium microdomains: organization and function. Cell Calcium 40: 405–412

Brown DA, Passmore GM (2010) Some new insights into the molecular mechanisms of pain perception. J Clin Invest 120: 1380–1383

Chaplan SR, Bach FW, Pogrel JW, Chung JM, Yaksh TL (1994) Quantitative assessment of tactile allodynia in the rat paw. J Neurosci Methods 53: 55–63

Choi SI, Hwang SW (2018) Depolarizing Effectors of Bradykinin Signaling in Nociceptor Excitation in Pain Perception. Biomol Ther 26: 255–267

Correll RN, Lynch JM, Schips TG, Prasad V, York AJ, Sargent MA, Brochet DXP, Ma J, Molkentin JD (2017) Mitsugumin 29 regulates t-tubule architecture in the failing heart. Sci Rep 7: 5328

Dray A, Perkins M (1993) Bradykinin and inflammatory pain. Trends Neurosci 16: 99–104

Gemes G, Bangaru ML, Wu HE, Tang Q, Weihrauch D, Koopmeiners AS, Cruikshank JM, Kwok WM, Hogan QH (2011) Store-operated Ca^2+^ entry in sensory neurons: functional role and the effect of painful nerve injury. J Neurosci 31: 3536–3549

Gerevich Z, Illes P (2004) P2Y receptors and pain transmission. Purinergic Signal 1: 3–10

Hadad N, Meyer HE, Varsanyi M, Fleischer S, Shoshan-Barmatz V (1999) Cardiac sarcalumenin: phosphorylation, comparison with the skeletal muscle sarcalumenin and modulation of ryanodine receptor. J Membr Biol 170: 39–49

Hirata Y, Brotto M, Weisleder N, Chu Y, Lin P, Zhao X, Thornton A, Komazaki S, Takeshima H, Ma J, Pan Z (2006) Uncoupling store-operated Ca^2+^ entry and altered Ca^2+^ release from sarcoplasmic reticulum through silencing of junctophilin genes. Biophys J 90: 4418–4427

Huang D, Huang S, Gao H, Liu Y, Qi J, Chen P, Wang C, Scragg JL, Vakurov A, Peers C, Du X, Zhang H, Gamper N (2016) Redox-Dependent Modulation of T-Type Ca^2+^ Channels in Sensory Neurons Contributes to Acute Anti-Nociceptive Effect of Substance P. Antioxid Redox Signal 25: 233–251

Ishikawa J, Ohga K, Yoshino T, Takezawa R, Ichikawa A, Kubota H, Yamada T (2003) A pyrazole derivative, YM-58483, potently inhibits store-operated sustained Ca^2+^ influx and IL-2 production in T lymphocytes. J Immunol 170: 4441–4449

Jin X, Shah S, Du X, Zhang H, Gamper N (2016) Activation of Ca^2+^-activated Cl^−^ channel ANO1 by localized Ca^2+^ signals. J Physiol 594: 19–30

Jin X, Shah S, Liu Y, Zhang H, Lees M, Fu Z, Lippiat JD, Beech DJ, Sivaprasadarao A, Baldwin SA, Zhang H, Gamper N (2013) Activation of the Cl^−^ Channel ANO1 by Localized Calcium Signals in Nociceptive Sensory Neurons Requires Coupling with the IP_3_ Receptor. Sci Signal 6:ra73

Kakizawa S, Kishimoto Y, Hashimoto K, Miyazaki T, Furutani K, Shimizu H, Fukaya M, Nishi M, Sakagami H, Ikeda A, Kondo H, Kano M, Watanabe M, Iino M, Takeshima H (2007) Junctophilin-mediated channel crosstalk essential for cerebellar synaptic plasticity. EMBO J 26: 1924–1933

Kirton HM, Pettinger L, Gamper N (2013) Transient overexpression of genes in neurons using nucleofection. Methods Mol Biol 998: 55–64

Kruger L, Kavookjian AM, Kumazawa T, Light AR, Mizumura K (2003) Nociceptor structural specialization in canine and rodent testicular "free" nerve endings. J Comp Neurol 463: 197–211

Lee B, Cho H, Jung J, Yang YD, Yang DJ, Oh U (2014) Anoctamin 1 contributes to inflammatory and nerve-injury induced hypersensitivity. Mol Pain 10: 5

Li H, Ding X, Lopez JR, Takeshima H, Ma J, Allen PD, Eltit JM (2010) Impaired Orai1-mediated resting Ca^2+^ entry reduces the cytosolic [Ca^2+^] and sarcoplasmic reticulum Ca^2+^ loading in quiescent junctophilin 1 knock-out myotubes. J Biol Chem 285: 39171–39179

Linley JE, Rose K, Ooi L, Gamper N (2010) Understanding inflammatory pain: ion channels contributing to acute and chronic nociception. Pflugers Arch 459: 657–669

Liu B, Linley JE, Du X, Zhang X, Ooi L, Zhang H, Gamper N (2010) The acute nociceptive signals induced by bradykinin in rat sensory neurons are mediated by inhibition of M-type K^+^ channels and activation of Ca^2+^-activated Cl^−^ channels. J Clin Invest 120: 1240–1252

Luo MC, Zhang DQ, Ma SW, Huang YY, Shuster SJ, Porreca F, Lai J (2005) An efficient intrathecal delivery of small interfering RNA to the spinal cord and peripheral neurons. Mol Pain 1: 29

Moriguchi S, Nishi M, Komazaki S, Sakagami H, Miyazaki T, Masumiya H, Saito SY, Watanabe M, Kondo H, Yawo H, Fukunaga K, Takeshima H (2006) Functional uncoupling between Ca^2+^ release and afterhyperpolarization in mutant hippocampal neurons lacking junctophilins. Proc Natl Acad Sci U S A 103: 10811–10816

Munoz F, Hu H (2016) The Role of Store-operated Calcium Channels in Pain. Adv Pharmacol 75: 139–151

Pacheco J, Ramirez-Jarquin JO, Vaca L (2016) Microdomains Associated to Lipid Rafts. Adv Ex Med Biol 898: 353–378

Parekh AB (2008) Ca^2+^ microdomains near plasma membrane Ca^2+^ channels: impact on cell function. J Physiol 586: 3043–3054

Petho G, Reeh PW (2012) Sensory and signaling mechanisms of bradykinin, eicosanoids, platelet-activating factor, and nitric oxide in peripheral nociceptors. Physiol Rev 92: 1699–1775

Qi Z, Wang Y, Zhou H, Liang N, Yang L, Liu L, Zhang W (2016) The Central Analgesic Mechanism of YM-58483 in Attenuating Neuropathic Pain in Rats. Cell Mol Neurobiol 36: 1035–1043

Qiu R, Lewis RS (2019) Structural features of STIM and Orai underlying store-operated calcium entry. Curr Opin Cell Biol 57: 90–98

Sahu G, Wazen RM, Colarusso P, Chen SRW, Zamponi GW, Turner RW (2019) Junctophilin Proteins Tether a Cav1-RyR2-KCa3.1 Tripartite Complex to Regulate Neuronal Excitability. Cell Rep 28: 2427–2442 e2426

Soderberg O, Gullberg M, Jarvius M, Ridderstrale K, Leuchowius KJ, Jarvius J, Wester K, Hydbring P, Bahram F, Larsson LG, Landegren U (2006) Direct observation of individual endogenous protein complexes in situ by proximity ligation. Nat Methods 3: 995–1000

Srikanth S, Jew M, Kim KD, Yee MK, Abramson J, Gwack Y (2012) Junctate is a Ca^2+^-sensing structural component of Orai1 and stromal interaction molecule 1 (STIM1). Proc Natl Acad Sci U S A 109: 8682–8687

Stefan CJ, Manford AG, Emr SD (2013) ER-PM connections: sites of information transfer and inter-organelle communication. Curr Opin Cell Biol 25: 434–442

Takeshima H, Hoshijima M, Song LS (2015) Ca^2+^ microdomains organized by junctophilins. Cell Calcium 58: 349–356

Taylor CW, Machaca K (2019) IP_3_ receptors and store-operated Ca^2+^ entry: a license to fill. Curr Opin Cell Biol 57: 1–7

Tian C, Du L, Zhou Y, Li M (2016) Store-operated CRAC channel inhibitors: opportunities and challenges. Future Med Chem 8: 817–832

Wei D, Mei Y, Xia J, Hu H (2017) Orai1 and Orai3 Mediate Store-Operated Calcium Entry Contributing to Neuronal Excitability in Dorsal Root Ganglion Neurons. Fron Cell Neurosci 11: 400

Weibrecht I, Leuchowius KJ, Clausson CM, Conze T, Jarvius M, Howell WM, Kamali-Moghaddam M, Soderberg O (2010) Proximity ligation assays: a recent addition to the proteomics toolbox. Expert Rev Proteomics 7: 401–409

Wolfrum C, Shi S, Jayaprakash KN, Jayaraman M, Wang G, Pandey RK, Rajeev KG, Nakayama T, Charrise K, Ndungo EM, Zimmermann T, Koteliansky V, Manoharan M, Stoffel M (2007) Mechanisms and optimization of in vivo delivery of lipophilic siRNAs. Nat Biotechnol 25: 1149–1157

Woo JS, Srikanth S, Nishi M, Ping P, Takeshima H, Gwack Y (2016) Junctophilin-4, a component of the endoplasmic reticulum-plasma membrane junctions, regulates Ca^2+^ dynamics in T cells. Proc Natl Acad Sci U S A 113: 2762–2767

Zhang F, Wang Y, Liu Y, Han H, Zhang D, Fan X, Du X, Gamper N, Zhang H (2019) Transcriptional Regulation of Voltage-Gated Sodium Channels Contributes to GM-CSF-Induced Pain. J Neurosci 39: 5222–5233

Zhang J, Carver CM, Choveau FS, Shapiro MS (2016) Clustering and Functional Coupling of Diverse Ion Channels and Signaling Proteins Revealed by Super-resolution STORM Microscopy in Neurons. Neuron 92: 461–478

